# Striking olfactory receptor gene repertoire expansion in Senegalese sole (*Solea senegalensis*)

**DOI:** 10.64898/2025.12.10.693368

**Authors:** Dorinda Torres-Sabino, Maialen Carballeda, Óscar Aramburu, Pablo Sánchez-Quinteiro, Diego Robledo, Carmen Bouza, Paulino Martínez

## Abstract

Chemoreception through olfaction is essential for regulating fish behaviour. Fish olfactory receptor repertoire comprises four multigene families (OlfC, OR, ORA and TAAR), whose diversity strongly shapes species-specific olfactory capabilities. In this study, the olfactory repertoire of the flatfish *Solea senegalensis* was characterized through orthology analysis with seven species, identifying 455 olfactory receptor genes in *S. senegalensis*, including large OlfC and TAAR gene expansions. Active functionality was shown in 426 genes of the total repertoire through the olfactory transcriptome and the RNA-seq libraries generated in this study, showing expression correlation within each family. Phylogenetic trees integrating orthologous and paralogous relationships, along with chromosomal locations were constructed. Notable synteny conservation was observed primarily among flatfish, partially lost with phylogenetically distant taxa. Olfactory receptor genes were heterogeneously distributed across twelve *S. senegalensis* chromosomes, constituting eleven major clusters of paralogous genes. Overall, *S. senegalensis* exhibits an expanded and specialized olfactory function, emerging as a promising model for studying chemoreception and addressing the reproductive issues in aquaculture.

## Introduction

Environmental perception modulates fish behaviour, which is fundamental for biological functions, such as feeding, reproduction or migration ^[1–5]^. As a result, fish possess a highly developed olfactory system, with some species exhibiting expanded and specialized olfactory capabilities that enhance their ability to detect chemical cues in the aquatic environment ^[6–8]^. Fish chemosensory detection through smell takes place at the olfactory organs, where each olfactory neuron expresses a single olfactory receptor belonging to one of four multigene families: olfactory receptors class C (OlfC/V2R in mammals), odorant receptors (OR), olfactory receptors class A (ORA/V1R in mammals), and trace amine-associated receptors (TAAR) ^[9–13]^.

The olfactory receptor gene repertoire shows large variation across vertebrate taxa, largely shaped by ecological adaptations ^[13–16]^. This diversity is primarily driven by rapid gene gain and loss events ^[17,18]^. However, the four olfactory receptor families in fish show distinct evolutionary dynamics ^[12,13]^. Particularly within teleost, the ORA family is small and evolutionary stable, comprising around six genes ^[19–20]^. On the contrary, the evolution of the OlfC, OR and TAAR families has experienced dramatic changes in their olfactory repertoire, with both important expansion and contraction events ^[12,13,21,22]^. In vertebrates, the olfactory receptor gene repertoire is primarily organized in genomic clusters and has largely expanded through tandem gene duplication events ^[8,11,16,17,23,24]^. These gene clusters may be further shaped by chromosomal rearrangements, including translocations^[7,25,26]^.

The increase of reliable genomic resources in fish in the last decade has greatly facilitated the study of their olfactory repertoire ^[13,27]^. A recent comprehensive phylogenetic analysis across 185 Actinopterygii species has shown great variation in their olfactory gene repertoire, ranging from 28 genes in a tetraodontiform species to 1,317 genes in a polypteriform species ^[12]^. Actinopterygii (ray-finned fish) constitute a large and diverse group within teleost. Among them, the order Pleuronectiformes, commonly known as flatfish, comprises over 600 species and has undergone rapid evolutionary divergence ^[28]^. Pleuronectiformes have successfully adapted to demersal habitats worldwide, characterized by limited light conditions and sediment-rich substrates, making olfaction an essential and specialized sensory organ for these species ^[29]^.

Within flatfish, Senegalese sole (*Solea senegalensis*) is an emerging European aquaculture species that faces a serious challenge related to the completion of its reproductive cycle in captivity ^[30,31]^. Reproductive issues have been hypothesized to stem from altered chemoreception abilities, potentially arising from differences at early life stages between captive rearing and natural environments ^[31–33]^. Despite its significance in the development of socio-sexual behaviours, knowledge of the genomic and functional basis of olfactory function in *S. senegalensis* remains limited. Recently, the full-length olfactory transcriptome of *S. senegalensis* has been characterized at the isoform level ^[34]^, which represents an essential resource to improve our knowledge of the olfactory gene receptor repertoire and deep into the potential reproductive failure of captive males.

In this study, a comparative genomic characterization of the olfactory repertoire of four Pleuronectiform species (*S. senegalensis*, *Cynoglossus semilaevis*, *Paralichthys olivaceus* and *Scophthalmus maximus*) was carried, including other three Teleostei with reliable genomic resources (*Astyanax mexicanus, Danio rerio and Oryzias latipes*), and a basal Holostei outgroup (*Lepisosteus oculatus*) to understand the evolutionary and functional diversification of the *S. senegalensis* olfactory repertoire. *S. senegalensis* showed to be the species with the largest olfactory repertoire among those studied, conformed by large gene clusters and isolated genes distributed across the genome originated through tandem gene duplications and translocation events. This information will contribute to future evolutionary and comparative analyses of olfactory systems in fish and vertebrates, with potential applications for *S. senegalensis* aquaculture.

## Results

### Olfactory gene receptor repertoire

The orthology analysis on the proteomes of the eight selected species retrieved the expected phylogeny (Supplementary Figure 1). Briefly, the four pleuronectiform species clustered together, and along with *O. latipes* formed the acanthopterygian cluster. The ostariophysians (*A*. *mexicanus* and *D*. *rerio*) appeared as a sister clade within teleost, and finally, the holostean *L*. *oculatus* was basal to teleost ^[28,35–37]^.

In total, 95 olfactory-related orthogroups were identified (19 OlfC, 57 OR, 5 ORA and 14 TAAR), comprising 1,868 genes across the eight species. The alignment of the CDS of the eight species against the olfactory receptor gene sequences by Policarpo et al. ^[12]^ corroborated most of the olfactory receptor genes (1,447 genes, 77.5%) identified through our orthology analysis and allowed retrieving six additional olfactory-related orthogroups containing a total of 26 genes. Therefore, the final dataset included 101 olfactory-related orthogroups (19 OlfC, 60 OR, 6 ORA and 16 TAAR) (Supplementary Table 1). Among these, 19 orthogroups contained genes from the eight species, whereas 28 were species-specific, with *D. rerio* and *L. oculatus* exhibiting the highest representation of single-species orthogroups.

Altogether, the curated dataset of olfactory gene repertoires comprised 1,894 genes across the eight species, distributed as follows: 446 OlfC, 936 OR, 41 ORA and 471 TAAR (Supplementary Table 1). Interestingly, *S*. *senegalensis* was the species with the largest olfactory gene repertoire, including 455 genes (195 OlfC, 125 OR, 6 ORA and 129 TAAR; Supplementary Table 2), followed by *D. rerio* (426), *A. mexicanus* (241) and *L. oculatus* (218). The remaining three flatfish species presented between 128 and 152 olfactory receptor genes (Figure 1).

**Figure 1.**
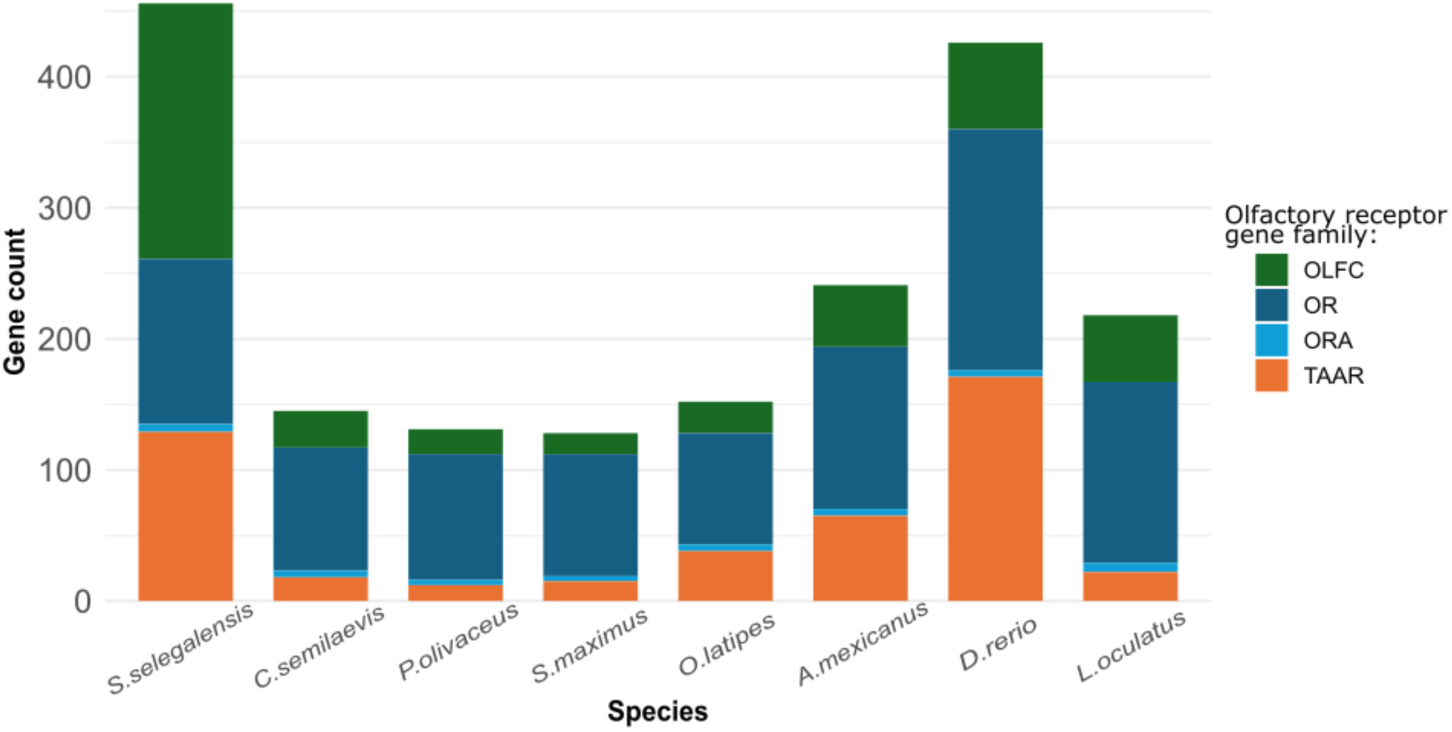
Stacked barplot of the number of genes in each of the four families of the olfactory gene repertoire of *Solea senegalensis*, *Cynoglossus semilaevis*, *Paralichthys olivaceus*, *Scophthalmus maximus*, *Oryzias latipes*, *Astyanax mexicanus*, *Danio rerio*, and *Lepisosteus oculatus*.

Overall, the analysis substantially enriched the known olfactory gene repertoires of six of the eight species, except for *A. mexicanus* and *L. oculatus*, as well as improved the available olfactory gene repertoire resources of *S. senegalensis*, highlighting a species with a large olfactory receptor repertoire (Table 1). Notably, all the 455 genes were protein coding genes that included functional domains directly related to chemoreception through olfaction.

**Table 1.**
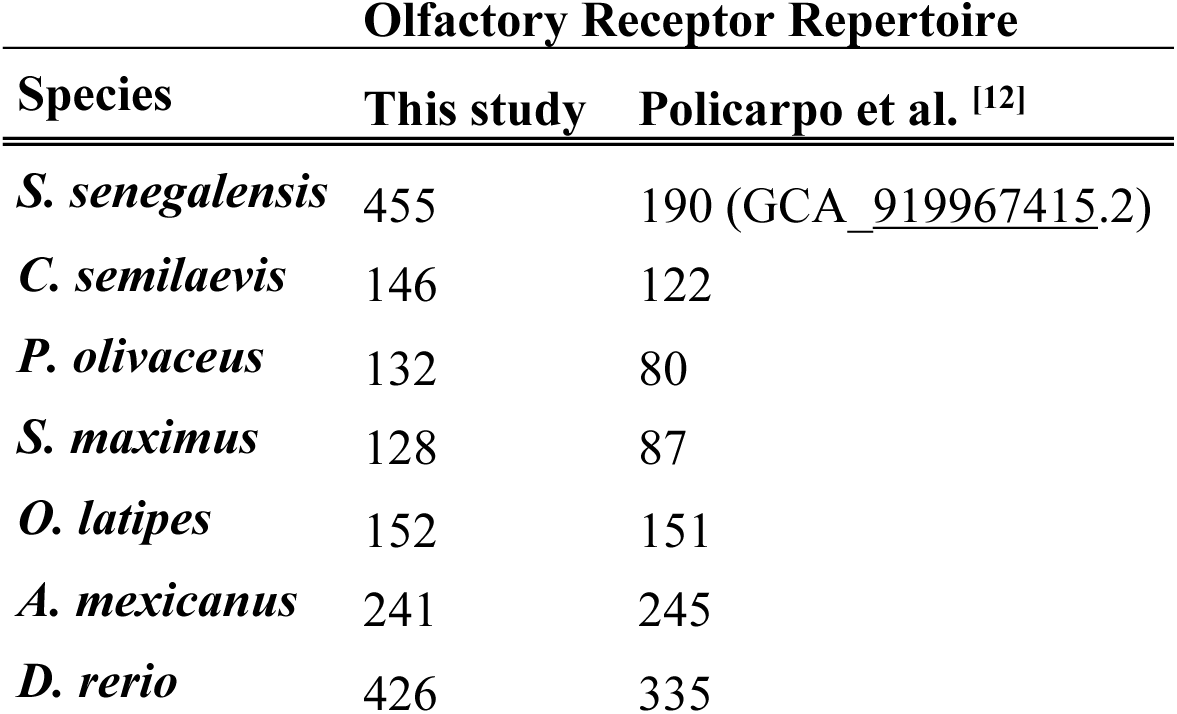
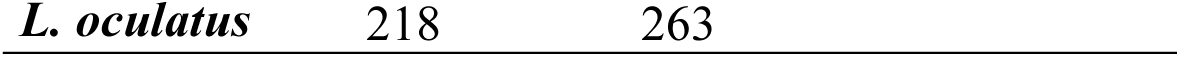
Comparison of olfactory receptor repertoires across eight species between our study and a previous study by Policarpo et al. ^[12]^. *Solea senegalensis* was not considered in the previous study, therefore its repertoire was compared with the reference genome annotations (GCA_919967415.2).

### *S. senegalensis* genome reannotation

In the *S. senegalensis* reference genome (GCA_919967415.2), 190 out of the 455 genes comprising the species olfactory gene repertoire were annotated as olfactory receptors. Notably, our search of non-annotated genes against Swissprot database resulted in 114,252 alignments. From these, a total of 3,597 genes (3,581 single-hits, 16 multiple-hits) were reannotated, representing 49.1% of the non-annotated Ensembl genes, 110 of them annotated as olfactory receptors. Moreover, this annotation categorized four genes as vomeronasal type 2 receptors (V2R), 5 as G protein-coupled receptors, 93 as calcium sensing receptors and four as taste receptor. The remaining 49 genes either lacked annotation or were associated with functions not directly related to olfaction (Supplementary Table 2).

### *S. senegalensis* olfactory receptor genomic landscape

Overall, olfactory receptor genes tended to cluster at specific genomic regions in the *S. senegalensis* genome. The physical map revealed large expansions, with genes mostly distributed across 12 chromosomes and a few ones to unassembled scaffolds (Figure 2). Eleven major olfactory receptor gene clusters mapped on seven chromosomes, comprising between 11 and 109 tandemly arranged genes belonging to a single olfactory receptor family. Meanwhile, six minor clusters were distributed across four chromosomes, and some single genes were scattered throughout the genome.

**Figure 2.**
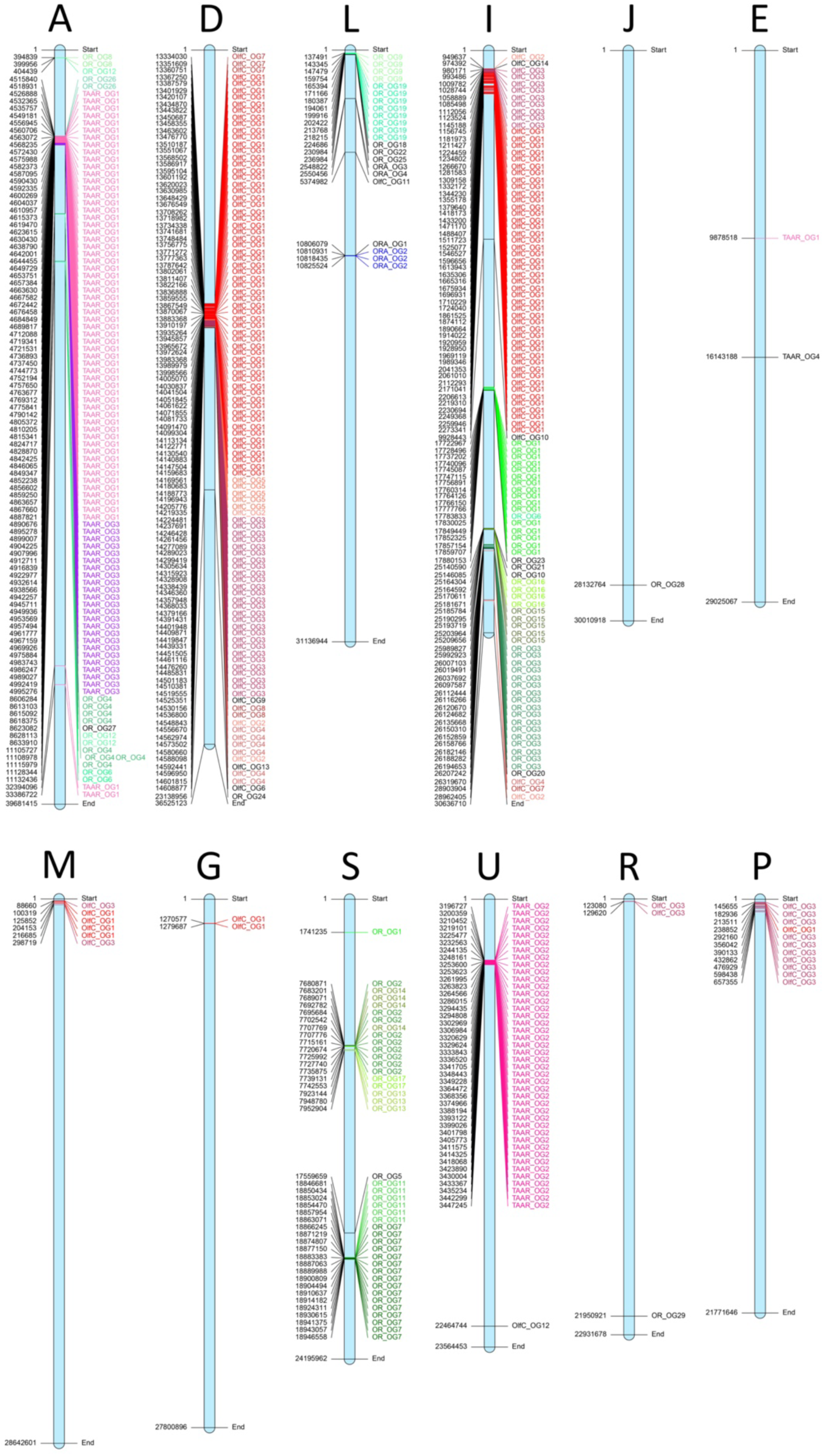
Physical map of the *Solea senegalensis* olfactory receptor genes within chromosomes. The name of the 12 olfactory gene bearing chromosomes is indicated at the top. The start and end coordinates of each chromosome is indicated, together with the start position of each gene (left side). The orthogroups for the different families are highlighted in different tones of red (OlfC), green (OR), blue (ORA) and pink (TAAR). In all cases, orthogroup number is indicated. Genes in black are isolated within an orthogroup.

Orthology analysis showed that the OlfC and TAAR families contain the largest *S. senegalensis* gene expansions, comprising 113 and 62 paralogous genes, respectively. Notably, the orthogroup OlfC-OG1 also included orthologs from all other analyzed species, although the total gene count varied among taxa. In contrast, TAAR-OG1 showed no representation in the non-pleuronectiform species studied except for *O. latipes*, which also pertains to the superorder Acanthopterigii. These observations suggest taxon and species-specific evolutionary trends. All *S. senegalensis* TAAR-OG1 genes, except one isolated gene on chromosome E, constituted a major cluster in chromosome A (Figure 3D). However, OlfC-OG1 appears to have undergone multiple tandem duplications and transposition or translocation events, reflected by the presence of two major clusters on chromosomes D and I, a minor cluster on chromosome M and several single genes on other chromosomes (Figure 3A). The OR orthogroups were broadly distributed across four chromosomes, forming both major and minor clusters in chromosomes S, L, I and A, along with some isolated genes (Figure 3B). Conversely, all ORA genes were confined to chromosome L, supporting the evolutionary stability of this family (Figure 3C).

**Figure 3.**
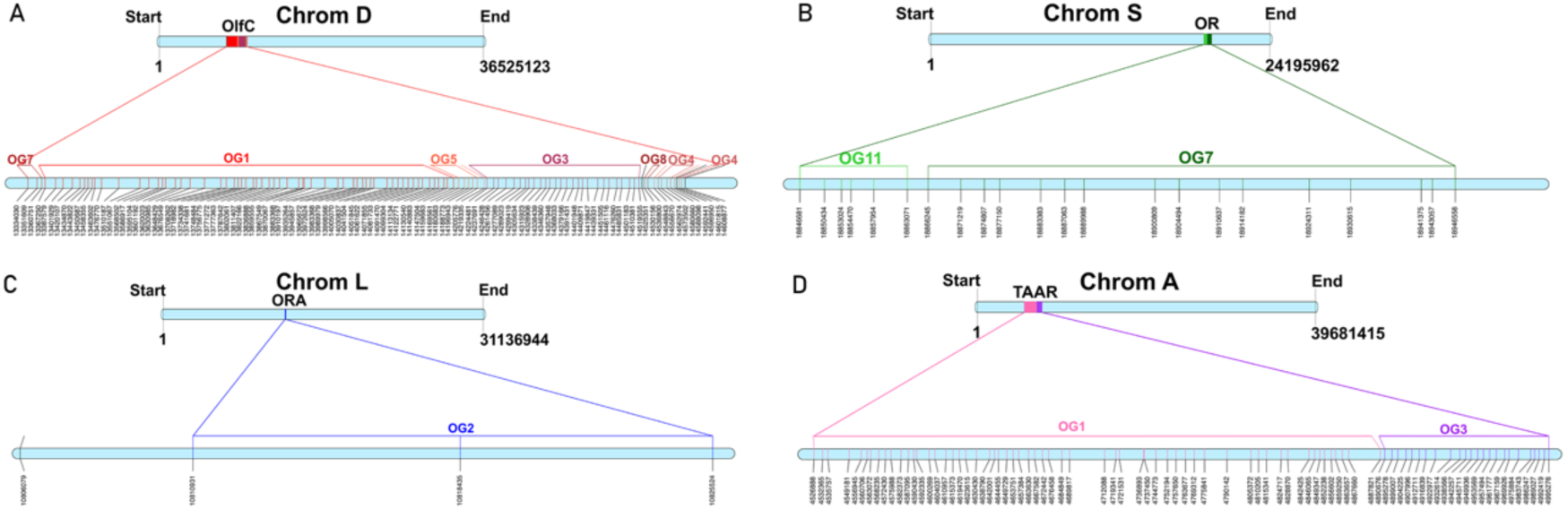
Diagram of the main clusters of olfactory receptor genes with their respective orthogroups (OG). The size of chromosomes and the start coordinate of each gene in bp are shown. Each vertical line within chromosomes correspond to a single gene. **(A)** OlfC major cluster in chromosome D; **(B)** Major OR cluster in chromosome S; **(C)** Minor ORA cluster and an isolated gene on chromosome L; **(D)** Major TAAR cluster in chromosome A.

### Olfactory receptor genes syntenic relationships across species

Globally, a substantial syntenic conservation of the olfactory receptor genes was observed across the eight species (Figure 4). In some cases, synteny was lost by the lack of orthologs, especially in the most distant species *A. mexicanus, D. rerio* and *L. oculatus*. Synteny was especially conserved among flatfish species and the closely related acanthopterygian *O. latipes*. Furthermore, some orthogroups split into more than one chromosome, supporting specific chromosomal rearrangements.

**Figure 4.**
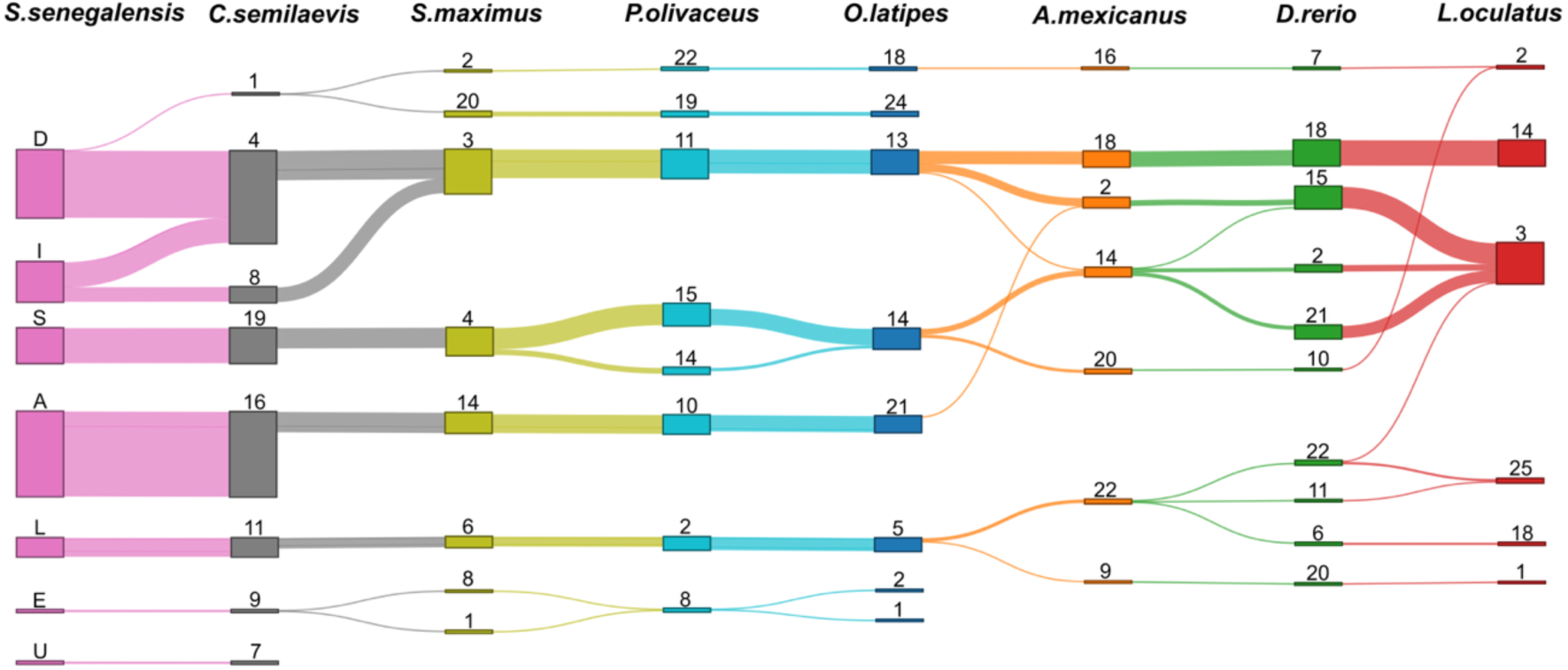
Syntenic relationships of the olfactory receptor gene-bearing chromosomes across the eight studied species. Chromosome identifiers (numbers/letters) are indicated, using a different colour per species. The line thickness represents the number of genes within syntenies across species from each chromosome.

### Olfactory receptor gene phylogeny

A phylogenetic tree was reconstructed for each of the four multigene olfactory receptor families OlfC, OR, ORA and TAAR (Figures 5, 6, 7 and 8, respectively). The basal genes suggested from *L. oculatus* phylogeny were used as the root for each of the multispecies tree (OlfC, OR and TAAR) that included the genes from *S. senegalensis*, *O. latipes* and *L. oculatus*. Conversely, the small ORA family allowed for the reconstruction of a phylogenetic tree for the eight species, in this case placing the root at the *L. oculatus* ORA1 gene, considered the basal gene of this family ^[38]^.

**Figure 5.**
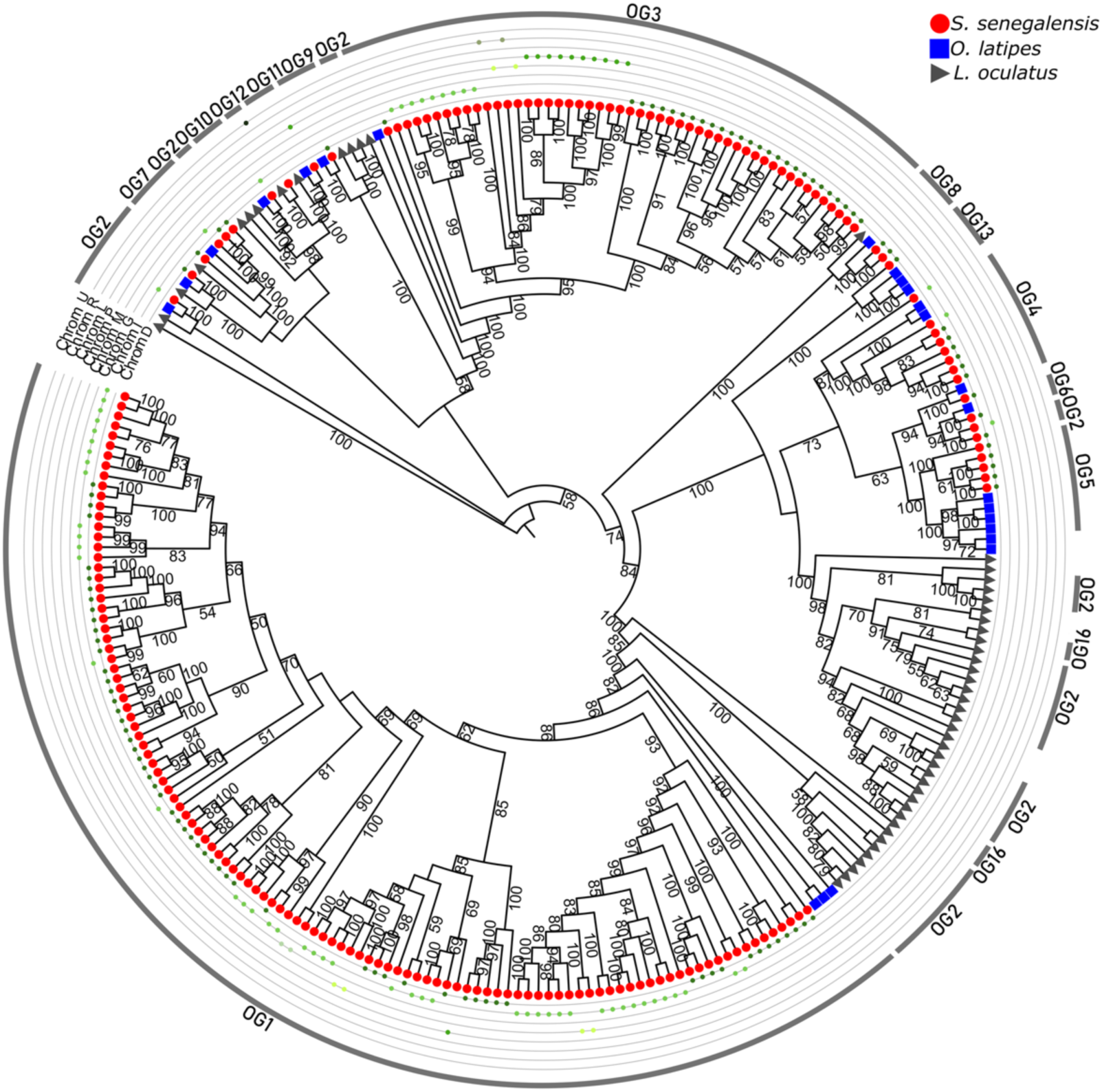
Phylogenetic tree of the OlfC olfactory receptor gene family using as outgroup the basal sequence of the L. oculatus tree for this family. Bootstrap values are shown at all nodes of the tree. Symbols at the end of each branch refer to each species analyzed. The next outer thin lines correspond to the *Solea senegalensis* chromosomes where gene clusters are located (ordered from outer to inner circles: U, L, R, P, M, G, I and D). The outer thick line shows the correspondence between clusters and the orthogroups (OG) where the genes are located.

The OlfC phylogeny was very robust, with most nodes showing 100% support and nearly all above 60%. After rooting using *L. oculatus* as outgroup, the tree split into two major clades. The first one included genes from the three species (*L. oculatus*, *O. latipes* and *S. senegalensis*), showing small expansions, altogether conforming a diverse clade, including different orthogroups, but also a large expansion of 52 *S. senegalensis* genes corresponding to a single orthogroup (OlfC-OG3). Despite being distributed across five *S*. *senegalensis* chromosomes, clustering was mainly observed in chromosomes D, I and P, which supports independent evolutionary events involving gene duplication and further translocation or transposition events. The other major node showed a basal branch with a few genes, and then, a great expansion into two main clades, the first one including a heterogeneous expansion of *L*. *oculatus*, constituted by different orthogroups, and the second one showing the largest and most recent *S. senegalensis* expansion, including 113 genes pertaining to OlfC-OG1. This expansion occurred in acanthopterygians, as shown by the basal clade including three *O. latipes* OlfC-OG1 genes, while the other large *S. senegalensis* OlfC expansions mainly mapped on chromosomes D and I (Figure 5).

The OR phylogeny was also highly confident with nearly all main nodes showing 100% bootstrap values. The evolutionary pattern of this family was sharply distinct to that of the OlfC family. From the root, the phylogenetic tree showed several consecutive small expansions of genes pertaining to different orthogroups of the three species studied, also involving *L*. *oculatus*. Then, a very confident node split into two main branches where species-specific clades of the three species appeared intermingled, supporting the established orthology and paralogy relationship retrieved from OrthoFinder. *S*. *senegalensis* OR genes clustered mainly in chromosomes A, I, L and S (Figure 6).

**Figure 6.**
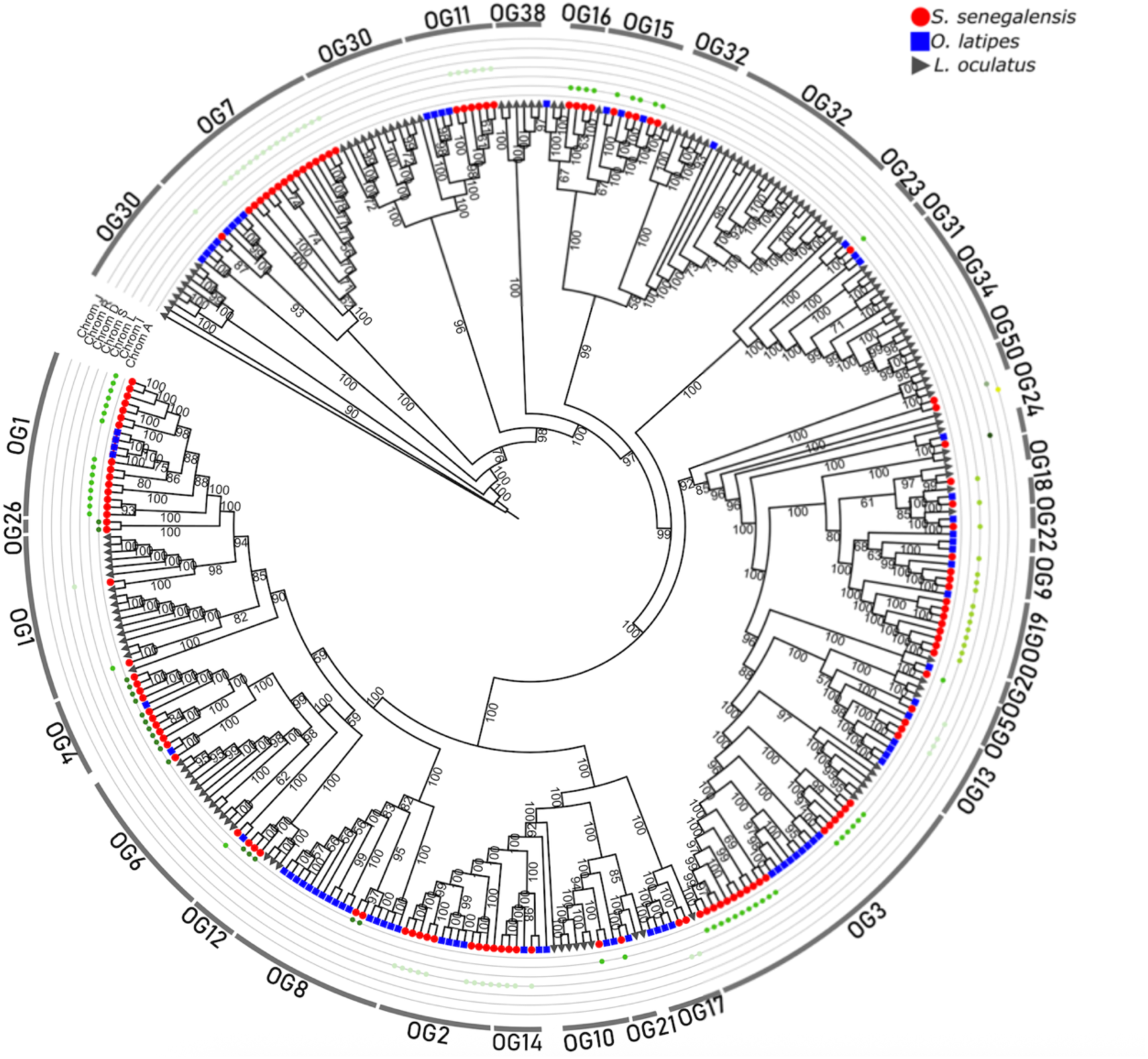
Phylogenetic tree of the OR olfactory receptor gene family using as outgroup the basal sequence of the *Lepisosteus oculatus* tree for this family. Bootstrap values are shown at all nodes of the tree. Symbols at the end of each branch refer to each species analyzed. The next outer thin lines correspond to the S. senegalensis chromosomes where gene clusters are located (ordered from outer to inner circles: J, R, D, S, L, I and A). The outer thick line shows the correspondence between clusters and the orthogroups (OG) where the genes are located.

The ORA phylogeny included the eight species and displayed a very robust tree including five confident clades that perfectly matched to the six orthogroups identified in our orthology analysis (Figure 7). Furthermore, all *S. senegalensis* ORA genes collocated on a single chromosome (L), in agreement with the comparative mapping and syntenic relationships observed across the species (Figure 4). *S. senegalensis* presented an expansion of three paralog genes in ORA-OG2.

**Figure 7.**
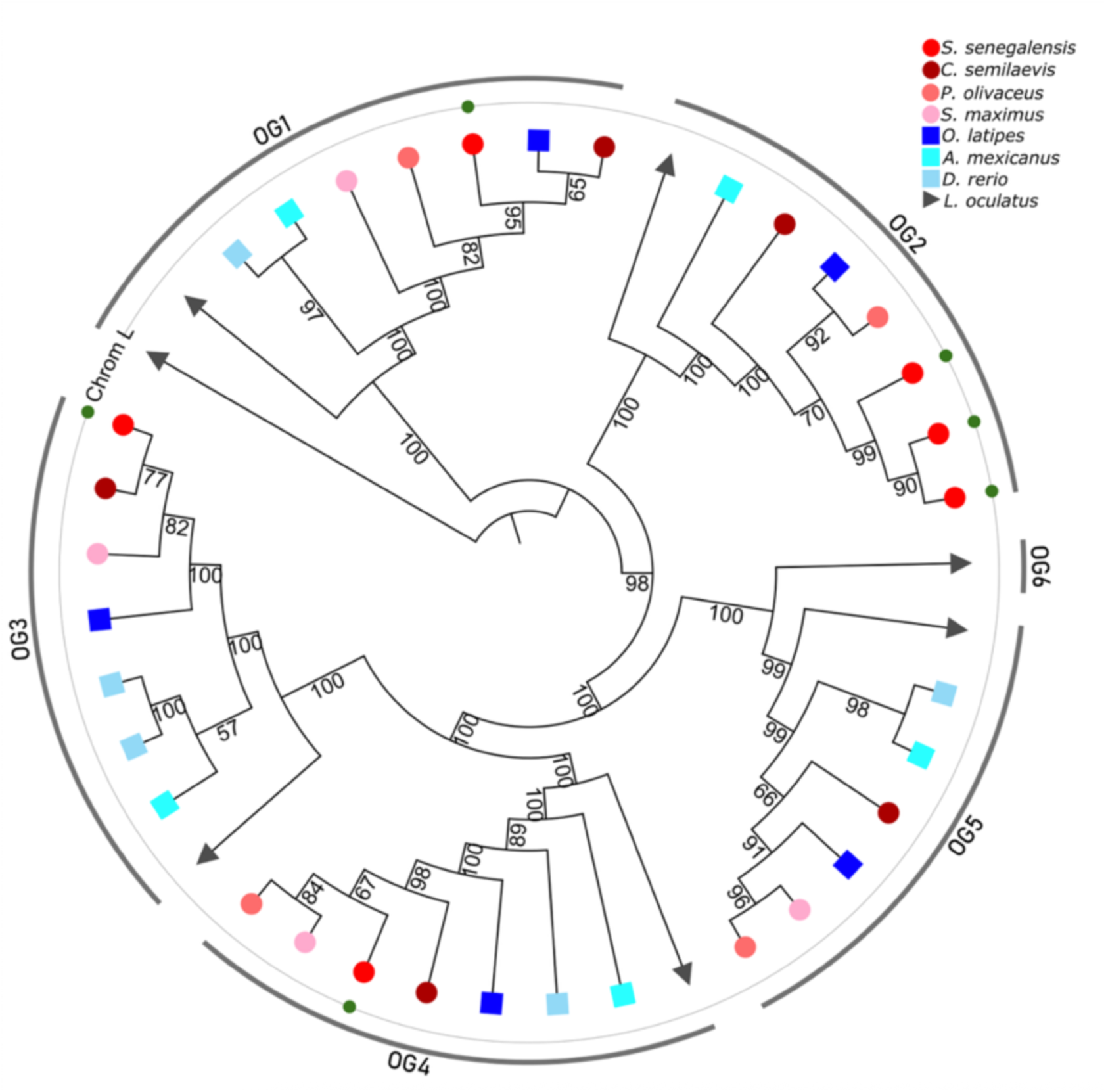
Phylogenetic tree of the ORA olfactory receptor gene family using as outgroup the basal sequence ORA1 family by Behrens et al. ^[38]^. Bootstrap values are shown at all nodes of the tree. Symbols at the end of each branch refers to each species analysed. The next outer thin line corresponds to the S. senegalensis chromosome where genes are located. The outer thick line shows the correspondence between clusters and orthogroups (OGs) are located.

The consistent TAAR family tree comprised 129 genes from *S. senegalensis*, resolved into two main highly confident nodes. These clades separate the ancestral lineage corresponding to *L. oculatus,* which first branched on a *L. oculatus* specific expansion. Then, the tree reflected the teleost genome duplication. Interestingly, a large species-specific gene expansion was observed for *S. senegalensis*, including 42 *S. senegalensis* genes corresponding to TAAR-OG2, intermingled with *O*. *latipes* orthologous genes included in the same orthogroup. Notably, all *S. senegalensis* genes from this cluster mapped on chromosome U, syntenic to the 13 *O. latipes* genes on chromosome 24 (Figure 4). The other main node split into clades including expansions of both teleost species. These genes pertained to TAAR-OG1, which split into one clade both for *O*. *latipes* and *S*. *senegalensis* orthologs, and another *S*. *senegalensis* specific clade of 50 genes pertaining to TAAR-OG1, tandemly arranged on chromosome A. This major gene cluster also included 24 *S. senegalensis* genes of TAAR-OG3 (Figure 8). These observations suggest that TAAR genes possibly emerged at different evolutionary times through tandem duplication.

**Figure 8.**
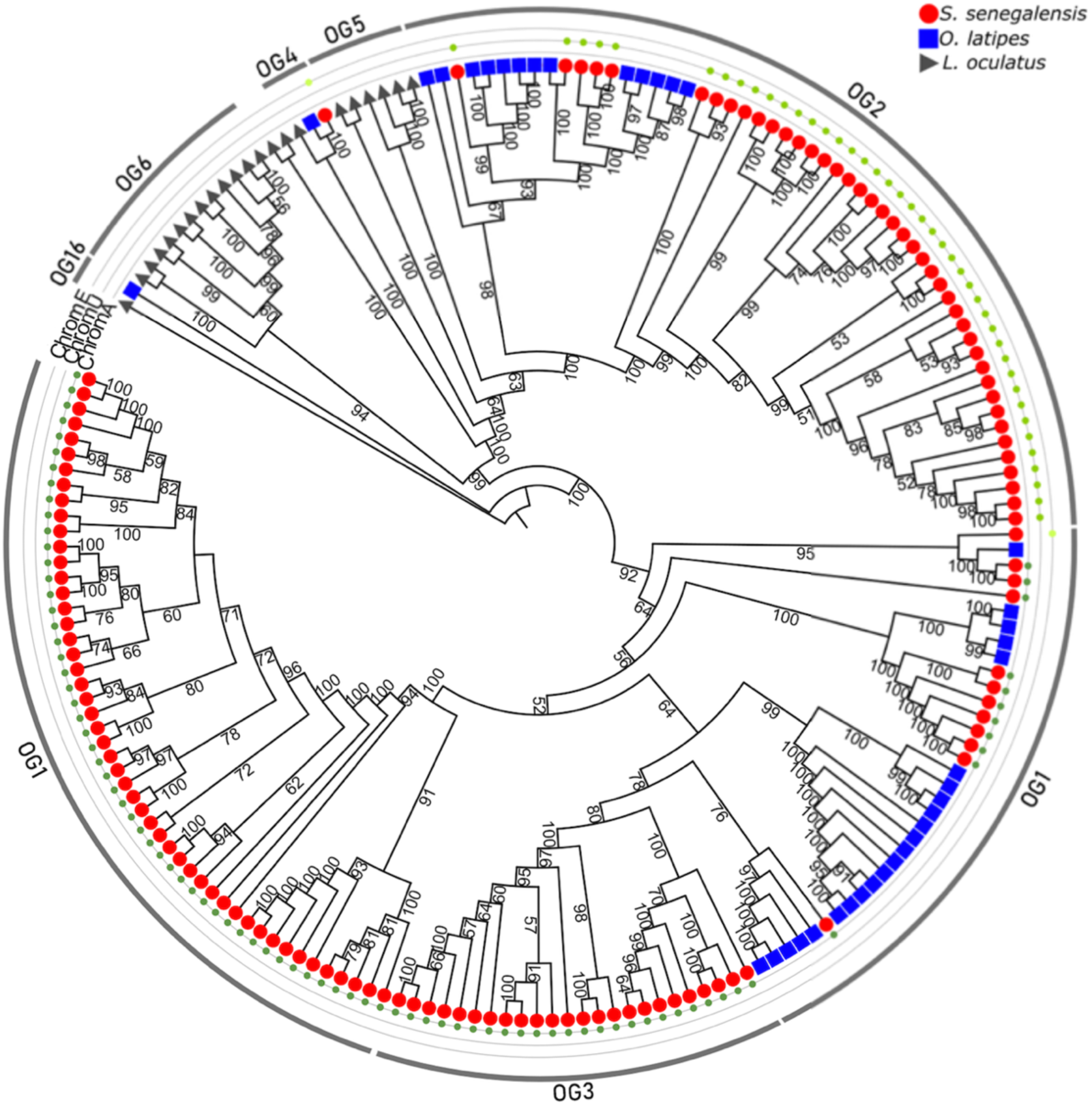
Phylogenetic tree of the TAAR olfactory receptor gene family using as outgroup the basal sequence of the *Lepisosteus oculatus* tree for this family. Bootstrap values are shown at all nodes of the tree. Symbols at the end of each branch refers to each species analysed. The next outer thin lines correspond to the *Solea senegalensis* chromosomes where gene clusters are located (ordered from outer to inner circles: E, U and A). The outer thick line shows the correspondence between clusters and orthogroups (OG) are located.

### *S. senegalensis* olfactory receptor gene expression

Among the reported olfactory repertoire in the present study, 321 genes were present in the olfactory transcriptome by Torres-Sabino et al. ^[34]^, and therefore showed active expression in the olfactory organ, meaning that over 70% of the predicted olfactory repertoire was found in the olfactory transcriptome dataset (Supplementary Figure 2). Since the olfactory transcriptome by Torres-Sabino et al. ^[34]^ was constructed based on a limited sample set, six RNA-seq olfactory organ libraries were generated in the present study to assess the expression of other genes, yielding on average 58.6 million (M) reads, after quality filtering, ranging from 50.3 to 71.5 M reads. On average, 90.5% of the reads mapped to the *S. senegalensis* assembly (range: 84.6% to 93.9%; Supplementary Table 3). A total of 389 olfactory receptor genes (171 OlfC, 115 OR, 5 ORA and 98 TAAR) were expressed with TPM ≥ 2, representing 85.5% of the total repertoire (Supplementary Table 4). Of these, 104 genes were inactive in the previously assembled olfactory transcriptome (Supplementary Figure 2), thus the olfactory repertoire increased up to 426 genes, leaving only 29 genes with no detectable expression.

Spearman correlation heatmap based on the expression of the olfactory receptor genes across the six RNA-seq libraries (Supplementary Table 4) suggested a certain clustering across the different olfactory receptor families, with most TAAR genes clustered in two groups, a high proportion of OR genes in a single group, and OlfC genes mainly in two clusters. Conversely, ORA genes were scattered across different groups (Figure 9A). Similarly, correlation networks using BioLayout showed a certain clustering per family (Figure 9B).

**Figure 9.**
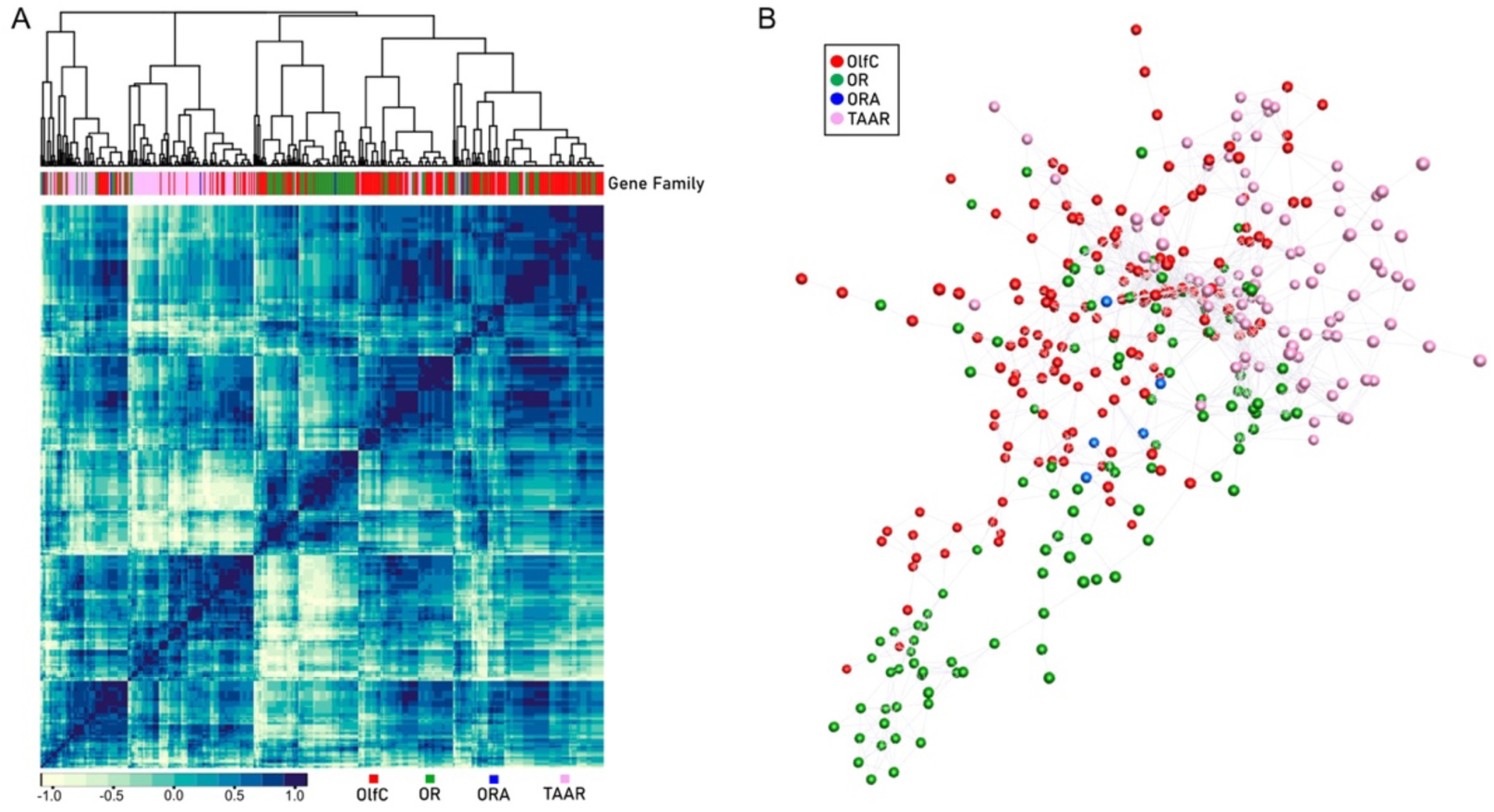
Expression correlation analysis across the four olfactory receptor gene families **(A)** Heatmap (Spearman correlation) of RNA-seq data on the olfactory receptor repertoires. **(B)** BioLayout correlation networks show how the expression of each olfactory receptor gene (dots) correlates across gene families. The diagram represents clustering on a network graph linking genes with a Pearson correlation coefficient (> 0.94), grouping genes with similar expression patterns. Clusters with less than six genes were removed.

## Discussion

In this study, the first comprehensive characterization of the olfactory gene repertoire of *S. senegalensis* enabled the identification of a remarkably large set of 455 olfactory receptor genes, representing the species with the largest olfactory repertoire within our study. The olfactory receptor repertoire previously annotated in the reference genome (GCA_919967415.2) included only 190 genes, highlighting the need to enhance olfactory receptor gene information across fish and vertebrate taxa.

### *S. senegalensis* olfactory receptor gene expansions

The large-scale phylogenetic analysis by Policarpo et al. ^[12]^ reported that the largest olfactory receptor gene family was generally OR, followed by TAAR and then OlfC. In contrast, *S. senegalensis* OlfC family was the largest, followed by TAAR and then OR, which may be related to species-specific functional adaptations. Indeed, the *S. senegalensis* OlfC family shows a large gene expansion of 113 paralogous genes located across five chromosomes, which showed conserved orthology relationships with the other seven species analyzed.

Previous studies reported the presence of lineage-specific OlfC gene expansions that conform clusters in other fish species, such as the 35 tandemly arranged genes in zebrafish ^[39]^, 55 in two distinct clusters in *Salmo salar* ^[23]^ and 61 in cichlid species ^[40]^. Thus, the OlfC cluster comprising 109 tandemly arrayed genes in *S. senegalensis* over 1.27 Mb represents to our knowledge the largest OlfC cluster reported in fish to date. The expansions of the OlfC multigene family, described as food-related amino acid receptors in teleost, may indicate a functional diversification in amino acid detection in *S. senegalensis*, as suggested in other teleost species ^[40,41]^. Furthermore, in ostariophysian fish, such as zebrafish, OlfC gene expansions via duplication enable finer detection of alarm pheromones from injured prey, driving fright reactions for predator avoidance ^[42]^. Similarly, in flatfish such as *S. senegalensis,* these receptors may perform analogous roles in sensing conspecific body fluids for threat detection and social behaviors.

In *S. senegalensis,* the TAAR family was the second most numerous, with three large expansions mainly located on two chromosomes. Lineage-specific TAAR expansions have been reported in other teleost species ^[12,43]^. Although their exact role in fish remains speculative, the detection of amine-related odors by TAAR receptors has been described to trigger aversion behaviours ^[44]^. A comparable situation is found in mammals, where TAAR receptors detect volatile amines that elicit strong behavioural reactions, including predator avoidance, social signaling and sex-dependent attraction ^[45]^.

Conversely, the 125 OR genes in *S. senegalensis* olfactory repertoire were dispersed across a large set of orthogroups, containing small subsets of paralogous genes. Although their physiological role in teleost species remains poorly understood, two OR zebrafish receptors have been involved in the detection of a specific prostaglandin that mediates mating behaviour ^[46]^. Recently, other studies have reported similar results in cichlids, pointing to the relevant role of OR receptors on sexual pheromone detection ^[47,48]^. The discovery of analogous receptors in *S. senegalensis* might offer insights into the poor mating performance of males on farms previously reported ^[31,33]^.

Finally, the six ORA genes identified in *S. senegalensis*, including the expansion of three paralogous genes, are collocated on chromosome L. Although little is known about their exact ligands in fish, there is evidence that ORA receptors detect bile acids and sexual pheromones depending on ORA ^[20,38,49]^, and particularly in zebrafish, the recognition of a pheromone has proved to trigger spawning behaviour ^[38]^.

Taken together, the advances in functional olfactory receptor studies focused on identifying chemical cues and elucidating their associated behavioural responses will offer valuable insights for improving reproductive protocols and behavioral management in aquaculture ^[50]^.

### Phylogenetic relationships and evolutionary pattern of olfactory receptor gene families

We explored orthology relationships together with available annotations in the species studied to characterize the olfactory repertoire and could identify 101 orthogroups. The comparison with previous large scale phylogenetic studies allowed us to validate most olfactory receptor genes and even enlarging the reported gene count in most species studied. Although the examined species come from diverse taxonomic groups, the analysis captured a complete orthology conservation on 19 olfactory receptor orthogroups, containing genes across the eight species. Furthermore, the identification of many paralog genes within these orthogroups, particularly in *S. senegalensis*, supports diverse evolutionary dynamics. This finding aligns with the theory that the olfactory receptor repertoires of teleost and other vertebrate taxa are shaped by species-specific adaptations driven by ecological and evolutionary pressures ^[12,14–16,18]^. Thus, the 28 species-specific orthogroups included a substantial portion of the olfactory receptor repertoire in the most evolutionarily distant species to *S*. *senegalensis*, to say *L. oculatus* and *D. rerio*.

The phylogenetic and syntenic analyses performed for each olfactory receptor gene families provided an integrated view of their evolutionary dynamics shaping *S. senegalensis* olfactory repertoire. Overall, the broad syntenic conservation across orthogroups of the four olfactory receptor families is concordant with the ancient origins of these families. Specifically, OR genes appear to constitute the ancient family, originated in chordates, whereas OlfC, ORA and TAAR would have emerged in the common ancestor of vertebrates ^[13]^. Conversely, the loss of synteny in other orthogroups, including some exhibiting *S. senegalensis* specific expansions, suggests the selection of diverse structural variants throughout their evolutionary history in response to environmental pressures. These patterns are reflected in the conserved ancestral OlfC clusters, with the large expansion in *S. senegalensis* OlfC-OG1 that emerged in acanthopterygians, mainly located on chromosome D. The heterogeneous OR clades distributed across several chromosomes is consistent with previous studies ^[9]^, and the strongly conserved ORA orthology relationships and synteny conservation is in accordance with previous works ^[20]^. Finally, the two TAAR clades support the diversification of the family through independent evolutionary events ^[21]^, with substantial synteny conservation between flatfish and *O. latipes*. All in all, these findings support that both conserved and lineage-specific evolutionary dynamics have shaped *S. senegalensis* olfactory repertoire, resulting in important differences with the other flatfish species analyzed as well as with the more distant teleost and holostean species.

### Adaptation of the olfactory function to demersal environments

Our study suggests a broadly diverse odorant detection in *S. senegalensis*, consistent with the size of its olfactory receptor repertoire ^[51,52]^. Its demersal lifestyle, characterized by low light irradiance and sediment-rich environment, has likely enhanced the sense of olfaction as an alternative to vision for environmental perception and intraspecific communication ^[28,29]^. A similar adaptation has been observed in the *A. mexicanus*, a blind species that inhabit caves, successfully adapted to dark environments, which exhibits enhanced olfactory sensitivity ^[53]^. Conversely, the much smaller repertoires of olfactory genes here described for the other flatfish analyzed in our study (*C. semilaevis*, *S. maximus* and *P. olivaceus)* and by Policarpo et al. ^[12]^, suggest a different adaptation strategy to demersal life, such as the specific visual-related or lateral line gene expansions reported in turbot ^[29]^ and tongue sole ^[54]^, respectively. Therefore, *S. senegalensis*, appears to follow an alternative sensory trajectory, where a markedly expanded olfactory repertoire may enhance the detection of trophic amino acids and other waterborne compounds released by buried or partially concealed prey, thereby improving foraging efficiency in environments where visual information is limited ^[8]^. It may also enhance spatial orientation and microhabitat recognition by exploiting the chemical heterogeneity of benthic environments. Furthermore, a broader olfactory receptor repertoire could facilitate intraspecific communication and reproductive signaling under visually restricted conditions ^[55]^.

## Conclusions

Although the genomic characterization and molecular basis of the olfactory response in teleost, particularly in flatfish, remain relatively unexplored, the extensive olfactory gene repertoire unveiled in *S. senegalensis* holds significant functional implications, underscoring the need for further investigation to elucidate the specific roles and mechanisms of these genes in the olfactory system of *S. senegalensis* and related flatfish species. Given that each receptor is thought to detect a specific array of ligands, including putative reproductive pheromones, elucidating their functional roles could offer insights for understanding how chemical cues regulate of fish social interactions and reproductive processes in fish. In this regard, these lineage-specific gene expansions offer a valuable opportunity to explore how olfactory repertoire diversification contributes to ecological specialization, demersal sensory adaptation, and potentially species-specific communication systems. Altogether, our findings position *S. senegalensis* not only as a valuable model for understanding olfactory evolution in flatfishes but also as a promising species in which to dissect the functional significance of chemosensory receptor diversity and the specific reproductive issues observed in farm captive males.

## Methods

### Comparative genomics of olfactory receptors

The proteomes of seven teleost species (*S. senegalensis* (GCA_919967415.2), *C*. *semilaevis* (GCA_000523025.1), *P*. *olivaceus* (GCA_024713975.2) *S*. *maximus* (GCA_013347765.1)*, O. latipes* (GCA_002234675.1)*, A*. *mexicanus (GCA_000372685.2)* and *D*. *rerio* (GCA_000002035.4)) and the outgroup *L. oculatus* (GCA_000242695.1) were downloaded from Ensembl (https://www.ensembl.org/) and beta Ensembl databases (https://beta.ensembl.org/, for *S. senegalensis* and *P. olivaceus*) in March 2025. A custom Python script was employed to only keep the protein sequence encoded by the coding DNA sequence (CDS) of each gene across the eight species, including the information of all exons.

Orthology relationships between species were explored with OrthoFinder v2.5.4 ^[56]^, applying default parameters. OrthoFinder reconstructs phylogenies from protein alignments and predicts gene function by inferring orthogroups. The analysis was performed at “Centro de Supercomputación de Galicia” (CESGA).

### Olfactory gene repertoire definition

Publicly available annotations of the studied species were retrieved from the Ensembl and NCBI (https://www.ncbi.nlm.nih.gov/) databases. We aimed to identify orthogroups that contained genes related to chemoreception through olfaction, included in one of the four fish olfactory receptor gene families (OlfC, OR, ORA and TAAR). If a gene within an orthogroup was consistently annotated as an olfactory receptor gene, functional homology was inferred across all the genes included in that orthogroup, and therefore, classified as olfactory receptor genes. Following this approach, a list of orthogroups containing olfactory receptor genes was retrieved, and the olfactory gene repertoires of each of the eight species included in our study were characterized.

The efficiency of the olfactory receptor gene repertoire characterization across the eight fish species was assessed by comparison against the repertoires reported across ray-finned species by Policarpo et al. ^[12]^. Briefly, the complete gene sequences available at https://figshare.com/articles/dataset/Olfactory_receptor_sequences_for_185_ray-finned_fishes/17061632 were downloaded and aligned with BLAST v2.13.0 ^[57]^ using blastn function (-outfmt 6, e-value 1e-20, max_target_seqs 1, other parameters by default) against the Ensembl CDS genome file of each of the eight species analyzed in the present study, retaining the best hit with e-value ≤ 1e-20. The resulting alignments were then compared by visual inspection to the olfactory gene repertoires characterized by orthology relationships in our study, thus confirming the common genes in both lists. Additional genes, not identified during our orthology analysis that could potentially be considered as olfactory receptor genes, were further examined.

### *S. senegalensis* genome reannotation

Annotation of 29,504 genes in the *S. senegalensis* genome was retrieved from beta Ensembl (GCA_919967415.2). Out of them, 7,326 genes lacked description or associated gene symbol, thus, their CDS sequences (8,621 sequences) were retrieved and aligned against swissprot_db’s protein database (UniProtKB) through BLAST v2.13.0 using blastx function (-outfmt 6, e-value 1e-5, num_threads 4, max_target_seqs 10, other parameters by default). The best alignment(s) were further filtered with a custom script by keeping that with the lowest e-value for each gene. In case of multiple alignments, further filtering was done to keep “single-hits” by applying hierarchical alignment criteria according to bit score, percent identity, and length of the alignment, in this order. Unsolved multiple alignments were tagged as “multiple-hits” but maintained for downstream analyses.

### Olfactory receptor gene repertoire: genomic landscape and phylogenetic analysis

The presence of olfactory receptor gene clusters across the *S. senegalensis* genome was inspected. A physical map of the genome showing the precise location of each olfactory receptor gene was generated using the R package LinkageMapView v2.1.2 ^[58]^. The chromosome name and size, gene start coordinates, olfactory receptor family and orthogroup were used as input. Minor olfactory clusters were defined as those containing three to ten tandemly arrayed genes, and major olfactory clusters as those comprising more than ten genes. For visual clarity, the physical map was further refined using Inkscape v.1.2 (https://inkscape.org). Additionally, the physical map was inspected by zooming in at major olfactory clusters for further examination of their gene composition and orthology relationships.

Syntenic relationships between *S. senegalensis* olfactory receptor repertoire regarding the other seven species were examined. The correspondence of each olfactory orthogroup across the eight species was established by investigating syntenic relationships based on the chromosomal locations of each olfactory receptor gene, considering only orthogroups with representation in *S. senegalensis*. Synteny blocks were represented with a Sankey plot using SankeyMATIC online tool (http://sankeymatic.com/build/).

A phylogenetic analysis was then conducted for each of the four multigene olfactory receptor families to investigate their evolutionary history by examining how orthogroup assignments corresponded to gene clustering across the genome. Given the considerable variation in total gene counts, different strategies were followed for phylogenetic reconstruction. The four olfactory receptor gene families were divided into two groups, one comprising the larger multigene families (OlfC, OR and TAAR), and another including the smaller ORA family. For the first group, the nucleotide sequences of the olfactory receptor genes from *S. senegalensis* were analysed together with *O. latipes*, as a well annotated acanthopterygian species, and *L. oculatus*, as a basal species before teleost genome duplication, used as outgroup. Rooting outgroup sequence for each family was chosen based on the consensus tree constructed for *L*. *oculatus* alone using the option -o of IQTREE2. Then, an alignment was performed for all genes from the three species pertaining to each gene family using MAFFT v7, with default parameters ^[59]^. Maximum-likelihood trees were constructed with IQ-TREE2 ^[60]^ using 1000 ultrafast bootstrap replicates ^[61]^ and optimal selection model defined by ModelFinder ^[62]^. For the small ORA family, an alignment for all the ORA genes identified in the eight species was performed using MAFFT, with default parameters, and the constructed phylogenetic tree was rooted on the ORA1 gene of *L*. *oculatus*, consistently established as the basal subfamily ^[38]^.

Bootstrap values > 50 were considered confident and included at each node of the trees. Around each tree, the chromosome location of each gene in the *S. senegalensis* genome was included, highlighting the correspondence of phylogenetic clades with gene clusters across the genome. Furthermore, the orthogroup genomic correspondence was indicated in cases where two or more genes belonging to the same orthogroup were closely clustered in the phylogenetic tree.

### Sample collection and RNA extraction

The olfactory organs of six 24 months old adult individuals were collected in June 2022 at Sea Eight SL (Safiestela, Portugal). All fish were maintained in indoor tanks at the standard water temperature and feeding conditions, following production protocols ^[63]^. Fish were sacrificed by decapitation under ethical regulations of the Sea Eight SL Company, in accordance with EU guidelines (96/609/EU). The six individuals were dissected, and samples were immediately flash frozen in liquid nitrogen before long-term storage at -80°C, until their use.

Total RNA was extracted using the E.Z.N.A. RNA Kit for Animal Tissue (Omega Bio-Tek, 2018, Product No. R6731). Briefly, frozen olfactory organs were homogenized using a TissueLyser (TissueLyser II, Qiagen) in GTC Lysis Buffer supplemented with 2-mercaptoethanol to preserve nucleic acids. A single stainless steel bead of 5 mm per tube was added for the homogenization process (1 min. 30,000 Hz; 1 min pause; 1 min 30,000 Hz). Afterwards, the lysates were centrifuged and passed through HiBind Mini Column, followed by RNA purification and elution with nuclease-free water.

### Library preparation and sequencing

RNA quantity and quality were assessed using the NanoDrop ND-1000 spectrophotometer (NanoDrop Technologies Inc.) and the Agilent 2100 Bioanalyzer System with RNA 600 Nano kit (Agilent Technologies). After successful quality controls, RNA samples were shipped to Novogene (Germany) for library preparation and sequencing on an Illumina NovaSeq 6000 S2 platform to generate 150 bp paired-end reads.

### Olfactory gene receptor expression validation

The expression of the *S. senegalensis* olfactory receptor gene repertoire identified in our study was evaluated through cross-checking with two different datasets: i) the number of olfactory receptor genes included in the reference full-length hybrid olfactory transcriptome ^[34]^, and ii) the olfactory receptor genes detected within the RNA-seq libraries generated in the present study. RNA-seq data were processed using the nf-core/rnaseq pipeline v3.10.1 ^[64]^ with default parameters. Briefly, FastQC was applied on the raw reads for quality assessment ^[65]^, and Trim Galore! trimmed adapter sequences and low-quality bases ^[66]^. Reads were aligned against the genome used as reference (GCA_919967415.2; by de la Herrán et al. ^[67]^) with STAR ^[68]^ and transcript read counts were normalized (transcripts per million; TPM) using RSEM ^[69]^. The expression threshold for transcriptionally active olfactory receptor genes was set to ≥ 2 TPM ^[70]^. Afterwards, the olfactory receptor gene expression overlapping across datasets was visualized using a Venn Diagram ^[71]^.

Spearman correlations were performed between the expression patterns of the four multigene families and represented using R package heatmap3 ^[72]^. Furthermore, olfactory receptor genes were clustered by expression using BioLayout ^[73]^ with a Pearson correlation threshold > 0.94, 2.1 cluster granularity and minimum of six genes per cluster.

## Limitations of the study

Although the present study provides promising foundations for a more comprehensive understanding of the fish olfactory system, revealing a highly developed and specialized olfactory function in the flatfish species *S. senegalensis*, the study possesses certain limitations regarding the functional validation of the data. Future studies could benefit from the findings here reported to generate functional data and behavioural consequences in the species, with potential direct applications in aquaculture production. Even though the present study reports the olfactory repertoire across the eight fish species, the olfactory receptor genes remain poorly annotated in public databases. Future studies focus on improving genomic annotations may enlarge the olfactory repertoires characterized in this study.

## Data and code availability

There is no original code reported in this study. Raw RNA-seq reads will be made available in a public repository upon acceptance of the manuscript.

## Acknowledgements

We thank the technical support and informatic resources provided by “Centro de Supercomputación de Galicia” (CESGA). This study was funded by: PID2022-137821OB-C31 funded by MCIN/AEI/10.13039/501100011033/FEDER, UE.; “Consellería de Cultura, Educación, Formación Profesional e Universidades Axencia Galega de Innovación” (“Xunta de Galicia”) (06_IN606D_2022_2693134; ED431C 2022/33). This study forms part of the Marine Science Programme (ThinkInAzul) supported by “Ministerio de Ciencia e Innovación” and “Xunta de Galicia” with funding from European Union NextGenerationEU (PRTR-C17.I1) and European Maritime and Fisheries Fund (2022-CP076).

## Author contributions

DTS, CB, DR and PM, conceptualization; DTS and PM, investigation; DTS and PM, methodology; DTS, MC, OA and PM formal analysis; DR and PM, funding acquisition; PM, project administration and supervision; DTS and PM, writing – original draft; DTS, MC, OA, PSQ, DR, CB and PM, review and editing.

## Supplementary files and legends

**Supplementary Figure 1.**
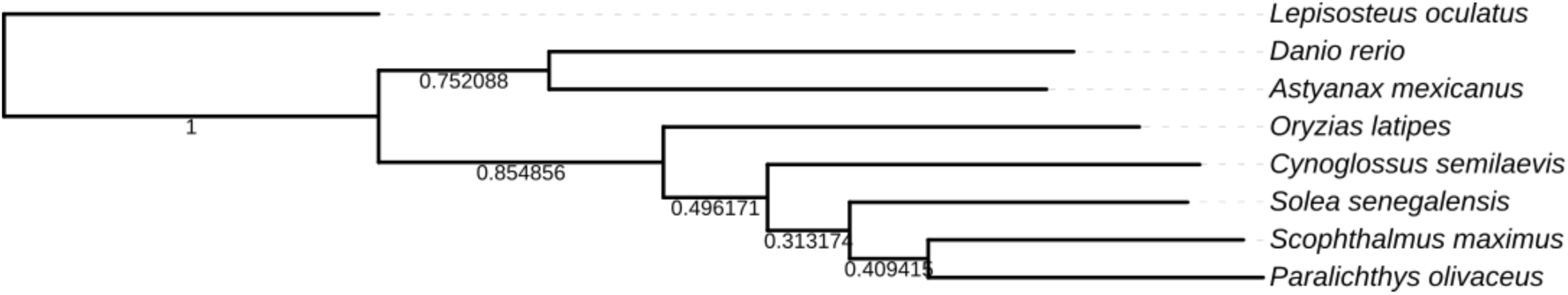
Phylogenetic tree of the eight species analyzed in this study, rooted in *Lepisosteus oculatus*. Branch lengths and the bootstrap values of each node are represented.

**Supplementary Figure 2.**
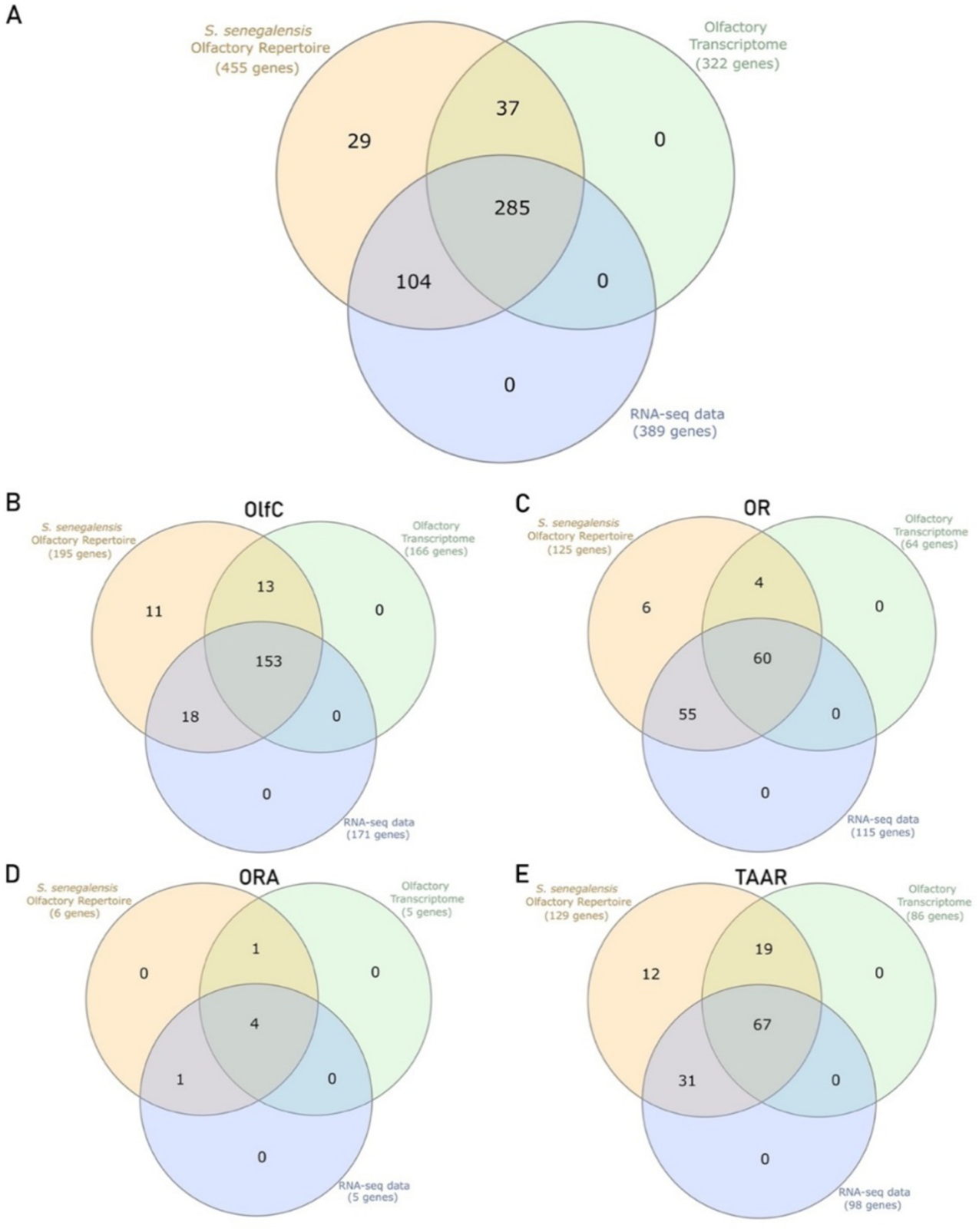
Venn Diagram comparing the datasets of the olfactory receptor repertoire (orange), the olfactory receptor genes expressed within the olfactory transcriptome (green) and the olfactory receptor genes expressed within our RNA-seq data (blue). **(A)** *Solea senegalensis* total olfactory gene repertoire. **(B)** OlfC genes. **(C)** OR genes. **(D)** ORA genes. **(E)** TAAR genes.

**Supplementary Table 1.** Olfactory receptor orthogroups of each of the four olfactory receptor gene families across the eight species.

**Supplementary Table 2.** *Solea senegalensis* olfactory receptor repertoire information.

**Supplementary Table 3.** Summary of the sequencing metrics of the six olfactory rosettes RNA-seq libraries, processed with the nf-core pipeline.

**Supplementary Table 4.** Expression (TPM) of the olfactory receptor genes in *Solea senegalensis* across six olfactory rosette samples.

## References

1. Hara T.J. (1975). Olfaction in fish. Prog Neurobiol. 5, 271–335. 10.1016/0301-0082(75)90014-3

2. Laberge, F., and Hara, T.J. (2001). Neurobiology of fish olfaction: a review. Brain Res. Rev. 36, 46–59. 10.1016/S0165-0173(01)00064-9

3. Hamdani, E.H., and Døving, K.B. (2007). The functional organization of the fish olfactory system. Prog. Neurobiol. 82, 80–86. 10.1016/j.pneurobio.2007.02.007

4. Kermen, F., Franco, L.M., Wyatt, C., and Yaksi, E. (2013). Neural circuits mediating olfactory-driven behavior in fish. Front. Neural Circuits 7, 62. 10.3389/fncir.2013.00062

5. Bowers, J., Li, C.-Y., Parker, C.G., Westbrook, M.E., and Juntti, S.A. (2023). Pheromone perception in fish: mechanisms and modulation by internal status. Integr. Comp. Biol. 63, 407–427. 10.1093/icb/icad049

6. Kasumyan, A.O. (2004). The olfactory system in fish: structure, function, and role in behavior. J. Ichthyol. 44, S180–S223.

7. Liu, H., Chen, C., Lv, M., Liu, N., Hu, Y., Zhang, H., Enbody E.D., Gao Z., Andersson L., and Wang W. (2021). A chromosome-level assembly of blunt snout bream (*Megalobrama amblycephala*) genome reveals an expansion of olfactory receptor genes in freshwater fish. Mol. Biol. Evol. 38, 4238–4251. 10.1093/molbev/msab152

8. Burguera, D., Dionigi, F., Kverková, K., Winkler, S., Brown, T., Pippel, M., Zhang Y., Shafer M., Nichols A.L.A., Myers E, et al. (2023). Expanded olfactory system in ray-finned fishes capable of terrestrial exploration. BMC Biol. 21, 163. 10.1186/s12915-023-01661-8

9. Alioto, T.S., and Ngai, J. (2005). The odorant receptor repertoire of teleost fish. BMC Genomics 6, 173. 10.1186/1471-2164-6-173

10. Saraiva, L.R., and Korsching, S.I. (2007). A novel olfactory receptor gene family in teleost fish. Genome Res. 17, 1448–1457. 10.1101/gr.6553207

11. Hussain, A., Saraiva, L.R., and Korsching, S.I. (2009). Positive Darwinian selection and the birth of an olfactory receptor clade in teleosts. Proc. Natl. Acad. Sci. USA 106, 4313–4318. 10.1073/pnas.0803229106

12. Policarpo, M., Bemis, K.E., Laurenti, P., Legendre, L., Sandoz, J.-C., Rétaux, S., and Casane, D. (2022). Coevolution of the olfactory organ and its receptor repertoire in ray-finned fishes. BMC Biol. 20, 195. 10.1186/s12915-022-01397-x

13. Korsching, S.I. (2025). Evolution of vertebrate olfactory receptor repertoires and their function. Curr. Opin. Behav. Sci. 61, 101483. 10.1016/j.cobeha.2025.101483

14. Vandewege, M.W., Mangum, S.F., Gabaldon, T., Castoe, T.A., Ray, D.A., and Hoffmann, F.G. (2016). Contrasting patterns of evolutionary diversification in the olfactory repertoires of reptile and bird genomes. Genome Biol. Evol. 8, 470–480. 10.1093/gbe/evw013

15. Policarpo, M., Baldwin, M.W., Casane, D., and Salzburger, W. (2024). Diversity and evolution of the vertebrate chemoreceptor gene repertoire. Nat. Commun. 15, 1421. 10.1038/s41467-024-45500-y

16. Kariyayama, H., Yusuke, O., Kashima, H., Nakanowatari, T., Harada, R., Yamaguchi, Y., and Suzuki, D.G. (2025). Hagfish olfactory repertoire illuminates lineage-specific diversification of olfaction in basal vertebrates. iScience, 10.1016/j.isci.2025.114118

17. Niimura, Y., and Nei, M. (2007). Extensive gains and losses of olfactory receptor genes in mammalian evolution. PLoS One 2, e708. 10.1371/journal.pone.0000708

18. Hughes, G.M., Boston, E.S.M., Finarelli, J.A., Murphy, W.J., Higgins, D.G., and Teeling, E.C. (2018). The birth and death of olfactory receptor gene families in mammalian niche adaptation. Mol. Biol. Evol. 35, 1390–1406. 10.1093/molbev/msy028

19. Zapilko, V., and Korsching, S.I. (2016). Tetrapod V1R-like ora genes in an early-diverging ray-finned fish species: the canonical six ora gene repertoire of teleost fish resulted from gene loss in a larger ancestral repertoire. BMC Genomics 17, 1–10. 10.1186/s12864-016-2399-6

20. Dong, X., Lv, M., Zeng, M., Chen, X., Wang, J., and Liang, X.F. (2025). Genome-wide identification and characterization of the ora (olfactory receptor class A) gene family and potential roles in bile acid and pheromone recognition in mandarin fish (*Siniperca chuatsi*). Cells 14, 189. 10.3390/cells14030189

21. Dieris, M., Kowatschew, D., and Korsching, S.I. (2021). Olfactory function in the trace amine-associated receptor family (TAARs) evolved twice independently. Sci. Rep. 11, 7807. 10.1038/s41598-021-87236-5

22. Zhang, Z., Sakuma, A., Kuraku, S., and Nikaido, M. (2022). Remarkable diversity of vomeronasal type 2 receptor (OlfC) genes of basal ray-finned fish and its evolutionary trajectory in jawed vertebrates. Sci. Rep. 12, 6455. 10.1038/s41598-022-10428-0

23. Johnstone, K.A., Lubieniecki, K.P., Koop, B.F., and Davidson, W.S. (2009). Genomic organization and evolution of the vomeronasal type 2 receptor-like (OlfC) gene clusters in Atlantic salmon, *Salmo salar*. Mol. Biol. Evol. 26, 1117–1125. 10.1093/molbev/msp027

24. Wang, Y., Sun, Y., and Joseph, P.V. (2023). Diverse evolutionary rates and gene duplication patterns among families of functional olfactory receptor genes in humans. PLoS One 18, e0282575. 10.1371/journal.pone.0282575

25. Wang, H., Wan, H.T., Wu, B., Jian, J., Ng, A.H., Chung, C.Y.L., Chow E.Y., Zhang J., Wong A.O.L., Lai K.P., et al. (2022). A chromosome-level assembly of the Japanese eel genome, insights into gene duplication and chromosomal reorganization. Gigascience 11, giac120. 10.1093/gigascience/giac120

26. Ye, M., Lin, X., Zhang, Y., Huang, Y., Li, G., and Tian, C. (2023). Genome-wide identification and characterization of olfactory receptor genes in silver sillago (*Sillago sihama*). Animals 13, 1232. 10.3390/ani13071232

27. Ahmad, S.F., Jehangir, M., Srikulnath, K., and Martins, C. (2022). Fish genomics and its impact on fundamental and applied research of vertebrate biology. Rev. Fish Biol. Fish 32, 357–385. 10.1007/s11160-021-09691-7

28. Lü, Z., Gong, L., Ren, Y., Chen, Y., Wang, Z., Liu, L., Li H., Chen X., Li Z., Luo H., et al. (2021). Large-scale sequencing of flatfish genomes provides insights into the polyphyletic origin of their specialized body plan. Nat. Genet. 53, 742–751. 10.1038/s41588-021-00836-9

29. Figueras, A., Robledo, D., Corvelo, A., Hermida, M., Pereiro, P., Rubiolo, J.A., Gómez-Garrido J., Carreté L., Bello X., Gut M., et al. (2016). Whole genome sequencing of turbot (*Scophthalmus maximus*; Pleuronectiformes): a fish adapted to demersal life. DNA Res. 23, 181–192. 10.1093/dnares/dsw007

30. Morais, S., Aragão, C., Cabrita, E., Conceição, L.E.C., Constenla, M., Costas, B., Dias, J., Duncan, N., Engrola, S., Estevez, A., et al. (2016). New developments and biological insights into the farming of Solea senegalensis reinforcing its aquaculture potential. Rev. Aquac. 8, 227–263. 10.1111/raq.12091

31. Martín, I., Riesco, M.F., Chaves-Pozo, E., Rodríguez, C., Martínez-Vázquez, J.M., Robles, V., Chereguini, O., and Rasines, I. (2021). Natural feed after weaning improves the reproductive status of *Solea senegalensis* breeders. Aquaculture 530, 735–740. 10.1016/j.aquaculture.2020.735740

32. Carazo Ortega, I. (2013). Comportamiento reproductivo y fisiología del lenguado senegalés (Solea senegalensis) en cautividad. PhD Thesis, Universitat de Barcelona.

33. Riesco, M.F., Valcarce, D.G., Martínez-Vázquez, J.M., Martín, I., Calderón-García, A.Á., González-Núñez, V., and Robles, V. (2019). Male reproductive dysfunction in *Solea senegalensis*: new insights into an unsolved question. Reprod. Fertil. Dev. 31, 1104–1115. 10.1071/RD18453

34. Torres-Sabino, D., Blanco, A., Villamayor, P.R., Rasines, I., Martín, I., Bouza, C., Robledo, D., and Martínez, P. (2025). Full-length hybrid transcriptome of the olfactory rosette in *Solea senegalensis*: an essential genomic resource for improving reproduction on farms. DNA Res. dsaf028. 10.1093/dnares/dsaf028

35. Pardo, B.G., Machordom, A., Foresti, F., Porto-Foresti, F., Azevedo, M.F.C., Bañón, R., Sánchez, L., and Martínez, P. (2005). Phylogenetic analysis of flatfish (Order Pleuronectiformes) based on mitochondrial 16S rDNA sequences. Sci. Mar. 69, 531–543. 10.3989/scimar.2005.69n4531

36. Betancur-R, R., Wiley, E.O., Arratia, G., Acero A., Bailly N., Miya M., Lecointre G., and Ortí G. (2017). Phylogenetic classification of bony fishes. BMC Evol. Biol. 17, 162. 10.1186/s12862-017-0958-3

37. Atta, C.J., Yuan, H., Li, C., Arcila, D., Betancur-R, R., Hughes, L.C., Ortí, G., and Tornabene, L. (2022). Exon-capture data and locus screening provide new insights into the phylogeny of flatfishes (Pleuronectoidei). Mol. Phylogenet. Evol. 166, 107315. 10.1016/j.ympev.2021.107315

38. Behrens, M., Frank, O., Rawel, H., Ahuja, G., Potting, C., Hofmann, T., Meyerhof, W., and Korsching, S. (2014). ORA1, a zebrafish olfactory receptor ancestral to all mammalian V1R genes, recognizes 4-hydroxyphenylacetic acid, a putative reproductive pheromone. J. Biol. Chem. 289, 19778–19788. 10.1074/jbc.M114.573162

39. Hashiguchi, Y., and Nishida, M. (2006). Evolution and origin of vomeronasal-type odorant receptor gene repertoire in fishes. BMC Evol. Biol. 6, 76. 10.1186/1471-2148-6-76

40. Nikaido, M., Suzuki, H., Toyoda, A., Fujiyama, A., Hagino-Yamagishi, K., Kocher, T.D., and Okada, N. (2013). Lineage-specific expansion of vomeronasal type 2 receptor-like (OlfC) genes in cichlids may contribute to diversification of amino acid detection systems. Genome Biol. Evol. 5, 711–722. 10.1093/gbe/evt041

41. Zhang, Z., Sakuma, A., Kuraku, S., and Nikaido, M. (2022). Remarkable diversity of vomeronasal type 2 receptor (OlfC) genes of basal ray-finned fish and its evolutionary trajectory in jawed vertebrates. Sci. Rep. 12, 6455. 10.1038/s41598-022-10428-0

42. Yang, L., Jiang, H., Wang, Y., Lei, Y., Chen, J., Sun, N., Zhao, H., and He, S. (2019). Expansion of vomeronasal receptor genes (OlfC) in the evolution of fright reaction in Ostariophysan fishes. Commun. Biol. 2, 235. 10.1038/s42003-019-0479-2

43. Hashiguchi, Y., and Nishida, M. (2007). Evolution of trace amine–associated receptor (TAAR) gene family in vertebrates: lineage-specific expansions and degradations. Mol. Biol. Evol. 24, 2099–2107. 10.1093/molbev/msm140

44. Hussain, A., Saraiva, L.R., Ferrero, D.M., Ahuja, G., Krishna, V.S., Liberles, S.D., and Korsching, S.I. (2013). High-affinity olfactory receptor for the death-associated odor cadaverine. Proc. Natl. Acad. Sci. 110, 19579–19584. 10.1073/pnas.1318596110

45. Zhang, J., Pacifico, R., Cawley, D., Feinstein, P., and Bozza, T. (2013). Ultrasensitive detection of amines by a trace amine-associated receptor. J. Neurosci. 33, 3228–3239. 10.1523/JNEUROSCI.4299-12.2013

46. Yabuki, Y., Koide, T., Miyasaka, N., Wakisaka, N., Masuda, M., Ohkura, M., Nakai, J., Tsuge, K., Tsuchiya, S., Sugimoto, Y., et al. (2016). Olfactory receptor for prostaglandin F2α mediates male fish courtship behavior. Nat. Neurosci. 19, 897–904. 10.1038/nn.4314

47. Li, C.-Y., Lawrence, K., Merlo-Coyne, J., and Juntti, S.A. (2023). Prostaglandin F2α drives female pheromone signaling in cichlids, revealing a basis for evolutionary divergence in olfactory signaling. Proc. Natl. Acad. Sci. USA 120, e2214418120. 10.1073/pnas.2214418120

48. Li, C.-Y., Bowers, J.M., Alexander, T.A., Behrens, K.A., Jackson, P., Amini, C.J., and Juntti, S.A. (2024). A pheromone receptor in cichlid fish mediates attraction to females but inhibits male parental care. Curr. Biol. 34, 3866–3880.e7. 10.1016/j.cub.2024.07.029

49. Cui, X., Chen, L., Tao, B., Zhang X., Song Y., Chen J., Duan M., Li W., Chen K., Pei Y., et al. (2025). Olfactory GnRH3 crypt sensory neurons transduce sex pheromone signals to induce male courtship behavior in zebrafish. Sci. China Life Sci. 68, 2191–2205. 10.1007/s11427-025-2917-5

50. Kamio, M., Yambe, H., and Fusetani, N. (2022). Chemical cues for intraspecific chemical communication and interspecific interactions in aquatic environments: applications for fisheries and aquaculture. Fish Sci. 88, 203–239. 10.1007/s12562-021-01563-0

51. Buck, L., and Axel, R. (1991). A novel multigene family may encode odorant receptors: a molecular basis for odor recognition. Cell 65, 175–187. 10.1016/0092-8674(91)90418-x

52. Miyasaka, N., Wanner, A.A., Li, J., Mack-Bucher, J., Genoud, C., Yoshihara, Y., and Friedrich, R.W. (2013). Functional development of the olfactory system in zebrafish. Mech. Dev. 130, 336–346. 10.1016/j.mod.2012.09.001

53. Blin, M., Valay, L., Kuratko, M., Pavie, M., and Rétaux, S. (2024). The evolution of olfactory sensitivity, preferences, and behavioral responses in Mexican cavefish is influenced by fish personality. eLife 12, e92861. 10.7554/eLife.92861.3

54. Chen, S., Zhang, G., Shao, C., Huang, Q., Liu, G., Zhang, P., Song, W., An, N., Chalopin, D., Volff, J.N., et al. (2014). Whole-genome sequence of a flatfish provides insights into ZW sex chromosome evolution and adaptation to a benthic lifestyle. Nat. Genet. 46, 253–260. 10.1038/ng.2890

55. Fatsini, E., Carazo, I., Chauvigné, F., Manchado, M., Cerdà, J., Hubbard, P.C., and Duncan, N.J. (2017). Olfactory sensitivity of the marine flatfish *Solea senegalensis* to conspecific body fluids. J. Exp. Biol. 220, 2057–2065. 10.1242/jeb.150318

56. Emms, D.M., and Kelly, S. (2019). OrthoFinder: phylogenetic orthology inference for comparative genomics. Genome Biol. 20, 238. 10.1186/s13059-019-1832-y

57. Camacho C, Madden T. BLAST+ Release Notes. 2013 Mar 12 [Updated 2025 Jul 22]. In: BLAST® Help [Internet]. Bethesda (MD): National Center for Biotechnology Information (US); 2008-. https://www.ncbi.nlm.nih.gov/books/NBK131777/

58. Ouellette, L.A., Reid, R.W., Blanchard, S.G., and Brouwer, C.R. (2018). LinkageMapView—rendering high-resolution linkage and QTL maps. Bioinformatics 34, 306–307. 10.1093/bioinformatics/btx576

59. Katoh, K., and Standley, D.M. (2013). MAFFT multiple sequence alignment software version 7: improvements in performance and usability. Mol. Biol. Evol. 30, 772–780. 10.1093/molbev/mst010

60. Minh, B.Q., Schmidt, H.A., Chernomor, O., Schrempf, D., Woodhams, M.D., von Haeseler, A., and Lanfear, R. (2020). IQ-TREE 2: new models and efficient methods for phylogenetic inference in the genomic era. Mol. Biol. Evol. 37, 1530–1534. 10.1093/molbev/msaa015

61. Hoang, D.T., Chernomor, O., von Haeseler, A., Minh, B.Q., and Vinh, L.S. (2018). UFBoot2: improving the ultrafast bootstrap approximation. Mol. Biol. Evol. 35, 518–522. 10.1093/molbev/msx281

62. Kalyaanamoorthy, S., Minh, B.Q., Wong, T.K.F., von Haeseler, A., and Jermiin, L.S. (2017). ModelFinder: fast model selection for accurate phylogenetic estimates. Nat. Methods 14, 587–589. 10.1038/nmeth.4285

63. Muñoz-Cueto, J.A., Mañanós, E.L., and Sánchez-Vázquez, E.J. (2019). The Biology of Sole. CRC Press 10.1201/9781315120393

64. Patel, H., Ewels, P., Peltzer, A., Botvinnik, O., Sturm, G., Moreno, D., et al. (2023). nf-core/rnaseq: nf-core/rnaseq v3.10.1 – Plastered Rhodium Rudolph. Zenodo. 10.5281/zenodo.7505987

65. Andrews S (2010). FastQC: A Quality Control Tool for High Throughput sequence Data. http://www.bioinformatics.babraham.ac.uk/projects/fastqc/

66. Martin, M. (2011). Cutadapt removes adapter sequences from high-throughput sequencing reads. EMBnet J. 17, 10–12. 10.14806/ej.17.1.200

67. de la Herrán, R., Hermida, M., Rubiolo, J.A., Gómez-Garrido, J., Cruz, F., Robles, F., Navajas-Pérez, R., Blanco, A., Villamayor, P.R., Torres, D., et al. (2023). A chromosome-level genome assembly enables the identification of the follicle-stimulating hormone receptor as the master sex-determining gene in the flatfish *Solea senegalensis*. Mol. Ecol. Resour. 23, 886–904. 10.1111/1755-0998.13750

68. Dobin, A., Davis, C.A., Schlesinger, F., Drenkow, J., Zaleski, C., Jha, S., Batut, P., Chaisson, M., and Gingeras, T.R. (2013). STAR: ultrafast universal RNA-seq aligner. Bioinformatics 29, 15–21. 10.1093/bioinformatics/bts635.

69. Li, B., and Dewey, C.N. (2011). RSEM: accurate transcript quantification from RNA-seq data with or without a reference genome. BMC Bioinformatics 12, 323. 10.1186/1471-2105-12-323

70. Wagner, G.P., Kin, K., and Lynch, V.J. (2013). A model based criterion for gene expression calls using RNA-seq data. Theory Biosci. 132, 159–164. 10.1007/s12064-013-0178-3

71. Heberle, H., Meirelles, G.V., da Silva, F.R., Telles, G.P., and Minghim, R. (2015). InteractiVenn: a web-based tool for the analysis of sets through Venn diagrams. BMC Bioinformatics 16, 169. 10.1186/s12859-015-0611-3

72. Zhao, S., Yin, L., Guo, Y., Sheng, Q., and Shyr, Y. (2021). heatmap3: an improved heatmap package. R package version 1.1.9. 10.1186/1471-2105-15-S10-P16

73 Theocharidis, A., van Dongen, S., Enright, A., and Freeman, T.C. (2009). Network visualization and analysis of gene expression data using BioLayout Express^3D^. Nat. Protoc. 4, 1535–1550. 10.1038/nprot.2009.177

